# Ferrochelatase regulates retinal neovascularization

**DOI:** 10.1101/2020.06.02.129650

**Authors:** Sardar Pasha Sheik Pran Babu, Darcy White, Timothy W. Corson

## Abstract

Ferrochelatase (FECH) is the terminal enzyme in heme biosynthesis. We previously showed that FECH is required for endothelial cell growth in vitro and choroidal neovascularization in vivo. But FECH has not been explored in retinal neovascularization, which underlies diseases like proliferative diabetic retinopathy and retinopathy of prematurity. Here, we investigated the inhibition of FECH using genetic and chemical approaches in the oxygen-induced retinopathy (OIR) mouse model. In OIR mice, FECH expression is upregulated and co-localized with neovascular tufts. Partial loss-of-function *Fech*^m1Pas^ mutant mice showed reduced retinal neovascularization and endothelial cell proliferation in OIR. An intravitreal injection of the FECH inhibitor *N*-methyl protoporphyrin had similar effects. Griseofulvin is an anti-fungal drug that inhibits FECH as an off-target effect. Strikingly, intravitreal griseofulvin blocked pathological tuft formation and revascularized areas of vasoobliteration faster than vehicle, suggesting potential as a FECH-targeting therapy. Ocular toxicity studies revealed that intravitreal injection of griseofulvin in adult mice does not disrupt retinal vasculature, function, or morphology. In sum, mutation and chemical inhibition of *Fech* reduces retinal neovascularization and promotes physiological angiogenesis, suggesting a dual effect on vascular repair upon FECH inhibition, without ocular toxicity. These findings suggest that FECH inhibitors could be repurposed to treat retinal neovascularization.

## INTRODUCTION

Neovascularization, the dysregulated growth of new blood vessels, plays a key role in a variety of sight-threatening disorders such as retinopathy of prematurity (ROP), proliferative diabetic retinopathy (PDR), and neovascular age-related macular degeneration (1, 2). Hypoxia and/or ischemia are strong promoters of this process. Vasculature defects resulting in hypoxia induce pathological new born blood vessels in ROP, which is a major cause of pediatric blindness (3, 4). The formation of new blood vessels is a multistep process which includes endothelial cell proliferation, migration, tubule formation and maturation. The endothelial cells and pericytes of the retinal circulation represent quiescent cell populations; however, they can re-enter the cell cycle due to vascular injury, cell loss or angiogenic stimuli (5). It has also been shown that vasorepulsive signals prevent vascular repair in highly damaged retina and brain (6, 7). Additionally, retinal neurons are affected by lack of blood supply. In pathological conditions, instead of vascular repair, formation of new blood vessels may occur (5), which is an abnormal growth of clusters of blood vessels that are fragile, can easily cause hemorrhage and do not lead to enough revascularization of ischemic retinal areas. The key pathological change, namely, retinal neovascularization, is associated with local ischemia followed by the subsequent development of neovascular tufts. The abnormal vascular changes and leakage may progress to retinal detachment. Inhibition of pathological angiogenesis represents a key therapeutic approach for ROP (8).

The crucial role of the vascular endothelial growth factor (VEGF) as a major regulator for most angiogenic functions of endothelial cells such as survival, hyper-permeability, migration, and proliferation has been well documented (9, 10) and thus anti-VEGF therapy for the eye has become a therapeutic mainstay (11). Indeed, the increasing use of intravitreal therapies targeting VEGF has transformed retinal disease care (12, 13). Anti-VEGF agents such as bevacizumab and ranibizumab show effective therapeutic potential either as monotherapy or in combination with laser therapy in ROP (14) and PDR patients (15, 16). However, since VEGF is essential for pro-survival signaling for blood vessel growth, treatment with anti-VEGF biologics could be associated with systemic and neurodevelopmental effects (17). Hence, there is a critical need for novel targets for treating pathological retinal neovascularization.

Previously, we identified ferrochelatase (FECH) as a target of an antiangiogenic natural product, cremastranone (18, 19). FECH is the final enzyme of the heme biosynthetic pathway, which catalyzes the insertion of ferrous iron into protoporphyrin IX (PPIX) to form heme. This pathway provides heme for hemoglobin and other vital hemoproteins (20–22). We showed that FECH mediates angiogenesis in vitro and in vivo (23). Interestingly, FECH can be inhibited by an FDA approved drug, the antifungal griseofulvin (24, 25), which alkylates the heme prosthetic group of cytochrome P450 enzymes to form the FECH inhibitor *N*-methylprotoporphyrin (NMPP) (26, 27). We showed that a single intravitreal injection of griseofulvin with or without combination with anti-VEGF drastically reduced choroidal angiogenesis in the laser-induced choroidal neovascularization mouse model (23). But the role of FECH and the therapeutic implications of its inhibition in retinal angiogenesis remains uncharacterized.

The mouse retina has been used extensively to study vascular pathologies and remodeling (8, 28–30). The oxygen-induced retinopathy (OIR) animal model enables study of vascular loss, regrowth, neovascular tufts, and regression, as seen in ROP in humans (31–34). Here, we investigated the inhibition of FECH using chemical and genetic approaches in the murine OIR model of retinal neovascularization. We show that FECH expression is upregulated and co-localized with neovascular tufts in the vascular pathology and *Fech* blockade reduced pathological retinal neovascularization and promoted physiological angiogenesis in the OIR model. Griseofulvin intravitreal treatment reduced neovascularization but did not lead to ocular toxicity, suggesting that this candidate might be repurposed as a treatment for neovascular eye disease.

## MATERIALS AND METHODS

### Animals and mouse strains

All mouse experiments were approved by the Institutional Animal Care and Use Committee, Indiana University School of Medicine and followed the guidelines of the Association for Research in Vision and Ophthalmology (ARVO) Statement for the Use of Animals in Ophthalmic and Visual Research. Wild-type C57BL/6J female mice, 6–8 weeks of age, plus timed E18-E19 stage pregnant mice were purchased from Jackson Laboratory (Bar Harbor, ME, USA). The mice were housed under standard conditions (35) in the Indiana University Laboratory Animal Resource Center (LARC). *Fech*^m1Pas^ mutant mice (36) were purchased from Jackson Laboratory on a BALB/c background and backcrossed into C57BL/6J; mixed-sex littermates from an F1 generation were used for experiments at 6–8 weeks of age. These animals were genotyped by qPCR according to a protocol from Jackson Laboratory (https://www2.jax.org/protocolsdb/f?p=116:5:0::NO:5:P5_MASTER_PROTOCOL_ID,P5_JRS_CODE:21675,002662). Sample sizes for experiments were based on power analyses, and treatments were randomly assigned by cage and animals. The experimenter was masked to genotype or treatment during analyses.

### Oxygen-induced retinopathy (OIR) mouse model

Mice were subjected to oxygen-induced retinopathy (OIR) as described previously (19, 30), (37). Briefly, postnatal day 7 (P7) mice, with their nursing mothers, were exposed to 75% O_2_(hyperoxia) in a hyperoxia chamber (Coy, Grass Lake, MI, USA) for 5 days (P7–P12) to initiate retinal vascular obliteration and returned to room air (normoxia) at P12. For temporal and spatial analysis of vascular pathology and FECH expression in the OIR mouse model, P12, P15 and P17 pups were euthanized at various time points. To monitor cell proliferation, S-phase marker and thymidine analogue, 5-ethynyl-2′-deoxyuridine, EdU (0.5 μg/g body weight) (Invitrogen, Waltham, MA, USA, A10044) was injected subcutaneously at different time points in OIR mice. To monitor tissue hypoxia, pimonidazole hydrochloride (60 mg/kg body weight) from Hypoxyprobe-Red549 Kit (Hypoxyprobe Inc, Burlington, MA, USA) was injected intraperitoneally at different time points in OIR mice.

### Intravitreal injections in juvenile and adult mice

Intravitreal injections were performed according to a previously published protocol (19, 38), (39). Mice were anesthetized for all procedures by intraperitoneal (i.p.) injections of 90 mg/kg ketamine hydrochloride and 5 mg/kg xylazine mixture. After intraocular injections, anesthesia was reversed with an i.p. injection of atipamezole hydrochloride (1 mg/kg). In OIR experiments, for P12 pups, the eyelid was opened using sterile tweezers to gain access to the globe. Pupils were dilated using 1% tropicamide and 2.5% phenylephrine. Proparacaine hydrochloride (0.5%) was used as topical anesthetic for the eyes. After dilation, a small incision was made at the nasal-temporal ora serrata (approximately 4 mm below the iris) to gain access to the posterior cavity (vitreous chamber) of the eye with the use of a 30-G insulin syringe needle. High precision, sterile Hamilton syringes (0.5-5 μL volume) with a sharp tip were used for injections. Vehicle (0.5 μL) with or without compound of interest was injected intravitreally without damaging the lens at the angle of about 45-60° towards the plane of injection. Successful injection led to mild perturbation in the anterior chamber; instant back flushing indicated a failed injection into a scleral pocket (e.g., between sclera and choroid) instead of injecting into the vitreous space.

In OIR experiments, at P12, after 2 hours of normoxia, pups were anesthetized using ketamine/xylazine. Based on the experiment, vehicle (DMSO), FECH inhibitor NMPP, or griseofulvin was intravitreally injected into each eye under a dissecting microscope as described above. After the injections and anesthesia reversal, pups along with the nursing mother were returned to normoxia (room air) conditions from P12 to P17. Both NMPP and griseofulvin dissolved readily in DMSO and were then diluted into phosphate-buffered saline solution to a final concentration of 0.5% DMSO. For mice, injecting 0.5 μL of a 100 μM aqueous solution of NMPP or griseofulvin, we estimated a final intravitreal concentration of 25 μM compound in young mice (based on the average vitreous volume of ~2 μL) and concentration of 10 μM compound in adult mice (based on the average vitreous volume of ~5 μL) (40, 41).

### In vivo optical coherence tomography (OCT) imaging

In vivo OCT was performed in griseofulvin injected adult mice as described previously (39, 42) at the indicated times using the Micron III intraocular imaging system (Phoenix Research Laboratories, Pleasanton, CA, USA). Briefly, before the procedure, mice were anesthetized with ketamine (90 mg/kg) and xylazine (5 mg/kg) and their pupils were dilated with 1% tropicamide solution (Alcon, Fort Worth, TX, USA) and lubricated with hypromellose ophthalmic demulcent solution (Gonak; Akorn, Lake Forest, IL, USA). Mice were then placed on a custom heated stage that moved freely to position the mouse eye for imaging. Several horizontal and vertical OCT images were taken in untouched, vehicle and griseofulvin-injected mice as indicated at various time points.

### Fluorescein angiography (FA)

Anesthetized mice were injected intraperitoneally with 25% fluorescein sodium (Fisher Scientific, Pittsburgh, PA, USA) at a dose of 50 μL per 25 g body weight (43). Fundus photography and FA were performed in untouched, vehicle-and griseofulvin-injected animals 14 days post intravitreal injection using the Micron III retinal imaging system and StreamPix software (Montreal, Quebec, Canada) at various times following fluorescein injection. The retinal vessels began filling about 30 seconds after fluorescein administration. Images of the central fundus were captured during early and late transit phases. Although timing varied due to variable rates of intraperitoneal absorption, retinal capillary washout usually occurred 5 minutes after dye administration.

### Electroretinogram (ERG) analysis

Full-field ERGs were recorded using a visual electrodiagnostic system (UTAS-E 2000; LKC Technologies, Gaithersburg, MD, USA) and analyzed as previously published (39). Briefly, mice were anesthetized as described above after dark adaptation overnight. Pupils were dilated and a drop of proparacaine hydrochloride (0.5%; Alcon) was applied to the cornea for topical anesthesia. ERG recordings were obtained simultaneously from both eyes with gold wire loop mouse corneal electrodes (STelesSR, LKC Technologies), with the reference electrode placed under the skin at the skull and the ground subdermal electrode at the tail. Flash ERGs were obtained from vehicle control and griseofulvin (100 μM) treated animals on day 10 post intravitreal injection. Briefly, scotopic rod recordings were performed on overnight dark-adapted mice, with 10 increasing light intensities of white light, and responses were recorded with a visual ERG stimulus presented at intensities of 0.025, 0.25, and 2.5 log cd∙s/m^2^ at 10-, 20-, and 30-second intervals, respectively. Ten responses were recorded and averaged at each light intensity. Photopic cone recordings were undertaken after mice were light adapted under the rod-saturating white background light of 100 cd∙s/m^2^ for 8–10 minutes. Recordings were performed with four increasing flash intensities from 0, 5, 10 and 25 log cd∙s/m^2^ in the presence of a constant 100 mcd∙s/m^2^ rod suppressing background light. The a-wave amplitude was measured from the baseline to the negative peak and the b-wave amplitude was measured from the a-wave trough to the maximum positive b-wave peak, behind the last prominent oscillatory potential. The average values of a- wave and b-wave amplitudes from scotopic 2.5 log cd∙s/m^2^ ERG (rod-driven response) and photopic 25 log cd∙s/m^2^ ERG (cone-driven response) were reported.

### Tissue preparation, cryosectioning, and immunohistochemistry

Whole eye cups without lens were fixed overnight at 4°C in 4% paraformaldehyde (PFA)/phosphate buffered saline (PBS) and cryoprotected in sucrose gradient (10-50%)/PBS solution overnight at 4°C. The samples were mounted in tissue embedding medium (Tissue-Tek). Frozen sections were cut at 15 μm on a cryostat (Leica Biosystems, Buffalo Grove, IL, USA), mounted on superfrost plus slides (Fisher Scientific, Pittsburgh, PA, USA) and stored at −80°C until use. Immunohistochemistry was performed using standard protocols (23, 39, 44). Cryosections were pretreated with PBS and DNase antigen retrieval, followed by 1 hour in blocking solution (5% bovine serum albumin (BSA) in 0.3% Triton-X 100-PBS). The primary antibodies were incubated in staining solution (0.5% BSA in 0.3% Triton-X 100-PBS) overnight. Primary antibodies included anti-FECH (rabbit, 1:250, LS Bio, Seattle, WA, USA, #LS-C409953), anti-IBA-1 (rabbit, 1:200, Wako, Richmond, VA, USA, #019-19741), anti-GFAP (2.2B10) (rat, 1:200, Invitrogen, #13-0300), and anti-cleaved Caspase 3(Asp175) (rabbit, 1:250, Cell Signaling, Danvers, MA, USA, #9661), plus isolectin GS-IB4 (Biotin, 1:250, Invitrogen, #121414). After incubation, cryoslides were washed 3×15 minutes in PBS. After washing, cryoslides were incubated for 2 hours at room temperature with secondary antibodies (AlexaFluor 488-, 555-, 647-, or Cy3-conjugated anti-rabbit, rat, mouse, goat IgG) (1:400, Life Technologies, Waltham, MA, USA, and Jackson Immunoresearch, West Grove, PA, USA), or, for isolectin GS-IB4, conjugated streptavidin DyLight 488 (1:200, Invitrogen). After the incubation, cryoslides were washed 3 times in PBS and then slides were fluoromounted using Vectashield mounting medium with DAPI (Vector Laboratories, Burlingame, CA, USA).

### Retinal flatmount preparation and whole mount staining

In OIR experiments, at various time points, pups were euthanized by decapitation, and the eyes were enucleated and fixed in 4% PFA for 2 hours at room temperature. The retinal flatmounts were prepared and immunostained as per a previously published protocol (45). Briefly, after removal of the optic nerve, the anterior eye parts and lens from the mouse eyes, neural retina was carefully dissected from the RPE– choroid– sclera complex and transferred to a glass slide. Four radial incisions were made ~2 mm apart from the site of the optic nerve with microdissection scissors to allow the retina to lie flat for imaging. In some cases, to prevent the curling of the retina and for straightening the outer edge of the retina, the periphery of the quadrants was removed using a no.11 scalpel blade. Then, retinal flatmounts were fixed again in 4% PFA overnight at 4°C. After fixation, retinal flatmounts were washed twice in PBS and then permeabilized for 2 hours in blocking buffer containing 0.5% Triton X-100 in 10% BSA prepared in PBS. After incubation, retinal flatmounts were stained over 2 to 3 days at 4°C in a shaker, for isolectin GS-IB4 (1:200) and FECH (1:200) in flatmount staining solution: 0.5% Triton X-100 in 1% BSA prepared in PBS. After the incubation, retinal flat mounts were washed 4×15 minutes in PBS and respective secondary antibodies were added to each retina in a dilution of 1:400 in diluted flatmount staining solution and incubated overnight at 4°C in a shaker protected from light.

In some cases, retinal flatmounts were combined with EdU and Hypoxyprobe-1 staining before or along with secondary antibody addition. EdU staining was performed in retinal flatmounts using Click-iT EdU Alexa Fluor 555 kits (Invitrogen) as per manufacturer instructions and hypoxic regions were detected in retinal flatmounts using anti-pimonidazole mouse DyLight 549 monoclonal IgG (Hypoxyprobe) following the manual provided. After staining, retinal flatmounts were washed 4×15 minutes in PBS. Finally, immunostained retinal flatmounts were transferred to a glass slide and cover-slipped with Fluoromount-G (Southern Biotechnology, Birmingham, AL, USA) and stored protected from light at 4°C.

### RNA extraction and cDNA synthesis

For RNA extraction, dissected retinas were either flash frozen (in liquid nitrogen) or collected in RLT (RNeasy kit, Qiagen, Germantown, MD, USA). Whole retinas were homogenized with a Qiashredder (Qiagen). Total RNA was extracted according to the manufacturer instructions (Qiagen RNeasy mini kit). The isolated RNA was eluted in 30 μL RNase-free water, quantified using a NanoDrop spectrophotometer and stored at – 80*°*C until further use. cDNA synthesis was performed with iScript cDNA synthesis kit (BioRad, Hercules, CA, USA) according to the manufacturer’s protocol and 0.5 μg RNA was used for the reverse transcription.

### Quantitative real time PCR (qPCR)

Quantification of mRNA was performed with the TaqMan Fast Advanced Master mix and TaqMan probes on a ViiA7 thermal cycler (Applied Biosystems, Foster City, CA, USA). PCR cycling conditions were as follows: 50°C for 2 minutes, 95°C for 10 minutes, 40 cycles at 95°C for 15 s and annealing temperature at 60°C for 1 minute 30 sec, followed by 95°C for 2 s. Gene expression was analyzed with the ViiA7™ Version 1.2 software. qPCR was performed in 10 μL volumes in a 384-well plate. Primer/probe sets used were as follows: *Vegfa* (Mm00437306_m1), *Fech* (Mm00500394_m1) and housekeeping controls *Hprt* (Hs03024075_m1) and *Tbp* (Mm01277042_m1). The data were analyzed using the ΔΔC_t_method. qPCR reactions were run with at least 3 biological replicates and as technical triplicates. The expression levels of genes were normalized to the two housekeeping genes and calibrated to the age matched, untouched sample.

### Protein preparation and immunoblot analysis

Retinas and choroids were snap frozen in liquid nitrogen immediately after dissection, and stored at −80°C. Pooled retina and choroid were lysed in 1×RIPA sample buffer (Thermo Scientific) containing 3% β-mercaptoethanol (Sigma, St. Louis, MO, USA), protease inhibitors (cOmplete mini, Roche, Indianapolis, IN, USA), and phosphatase inhibitors (PhosSTOP, Roche) on ice for 20 minutes, homogenized, and clarified by centrifugation at 12,000 × *g* for 15 min at 4°C. Supernatants were collected and protein concentration was determined using a Bradford assay. Equal amounts of total protein (30–50 μg) from each sample were resolved by 4–20% Tris-Glycine gels (nUview Precast gel, NB10-420, NuSeP, Germantown, MD, USA) and then transferred onto polyvinylidene fluoride membranes (Millipore, Burlington, MA, USA). Proteins were immunoblotted with antibodies against FECH (rabbit, 1:400, LS Bio, #LSC409953, and β-actin (mouse, 1:5000, Sigma-Aldrich, AC40). Secondary antibodies, anti-rabbit IgG peroxidase conjugated (1:10,000, Rockland, Rockland, MD, USA, #39116) and anti-mouse IgG peroxidase conjugated (1:10,000, Rockland, #37192) were used. All the dilutions were made in Tris Buffered Saline-0.05% (v/v) Tween-20 buffer containing 2% (w/v) bovine serum albumin (BSA). Immunoreactive bands were detected using Amersham ECL prime immunoblotting detection reagents (GE Healthcare, Chicago, IL, USA) on an Azure c600 Chemiluminescent imager (Azure Biosystems, Dublin, CA, USA).

### H & E paraffin staining and analysis

Eyes were enucleated and fixed in 4% PFA overnight, and then the eyes were paraffin embedded and sectioned at 5 μm thickness by the Indiana University School of Medicine Histology Core. Mayer’s hematoxylin and eosin staining was performed as described previously (39, 46). Retinal thickness quantified by calculating the ratio of A (the distance from the ganglion cell layer (GCL) to the outer edge of the inner nuclear layer (INL)) to B (the distance from the GCL to the outer edge of the outer nuclear layer (ONL)) and retinal morphology was quantified by layer wise cell nuclei counts across the vertical columns in the retinal sections (47).

### Image analysis and Quantification

For retinal flatmount overview image acquisition, tile scan images were acquired on an LSM700 confocal microscope (Plan-Apochromat 10× objective; Zeiss, White Plains, NY, USA) and stitched using Zeiss Zen black or blue software. For H & E stained retinal overview images, approximately 10 images were taken for each retina on an EVOS XL digital imaging system (4× objective) and all these images were stitched using photo merge automation in Photoshop CS7 (Adobe, Mountain View, CA, USA). Retinal vasculature area, vasoobliteration and neovascularization area were quantified as described previously (Supplemental Fig. S5) (30, 48). For EdU data analysis, images were acquired from the central region of interest in retinal flat mounts at least 1 mm away from the optic nerve head (center of the retina) and from at least four different central regions per retina (Supplemental Figs. S6 and S8). Cell counts were normalized to 1 mm^2^ retina area. Optical Z-sectioning was performed at 1 μm intervals for cell quantification and co-localization of double-positive cells using a Plan-Apochromat 20× objective on the Zeiss confocal microscope. Quantitative analyses were performed on flattened image stacks processed with Zeiss Zen Blue software. All images were cropped, and minor contrast and brightness adjustments were made in Adobe Photoshop CS7 and aligned with line drawings in Adobe Illustrator CC 2018.

### Statistical analysis

The data obtained from all experiments are represented as mean ± SEM unless otherwise indicated and were analyzed by either unpaired, two-tailed Student’s t-test or one-way ANOVA with Dunnett’s post hoc tests for comparisons between compound treatments and control. All analyses were performed using Prism software (v. 8.0; GraphPad, San Diego, CA, USA). A P value of < 0.05 was considered statistically significant in all tests.

## RESULTS

### Hypoxia-driven vascular pathology upregulates FECH expression in OIR

To investigate the temporal and spatial relationship between ischemic vascular pathology and FECH expression, we examined the OIR murine retina at different time points (**Fig. 1*A***). First, we sought to determine retinal vascular pathology and retinal hypoxia in OIR mice using vasculature and pimonidazole adduct staining. Temporal analysis of vascular staining using *Griffonia simplicifolia* isolectin B4 (GS-IB4) from P12 to P17 in OIR retina showed the expected biphasic vascular pathology: an initial vasoobliteration phase and a subsequent pathological neovascularization phase as previously described (30, 48, 49). Pimonidazole adduct immunostaining revealed substantial hypoxia in the central avascular regions and diffuse patterns in the peripheral vascular regions in OIR (Fig. 1*B*) compared to the control retina. Further, temporal analysis in OIR retina showed increased immunoreactivity for the glial fibrillary acidic protein (GFAP), suggesting Müller glia and astrocyte activation in response to retinal disease (50) and very little neuronal and vascular cell death (as evidenced by activated caspase-3 staining) in the OIR mouse model (Supplemental Fig. S1). Further, mRNA analysis showed a significant upregulation of *Vegfa* expression even after 2 hours of hypoxia at P12 and expression was persistent until P17 in OIR retina, consistent with previous studies (51, 52). Also, significant upregulation of *Fech* mRNA was observed at P17 in OIR murine retina (Fig. 1*C*).

**Figure 1.**
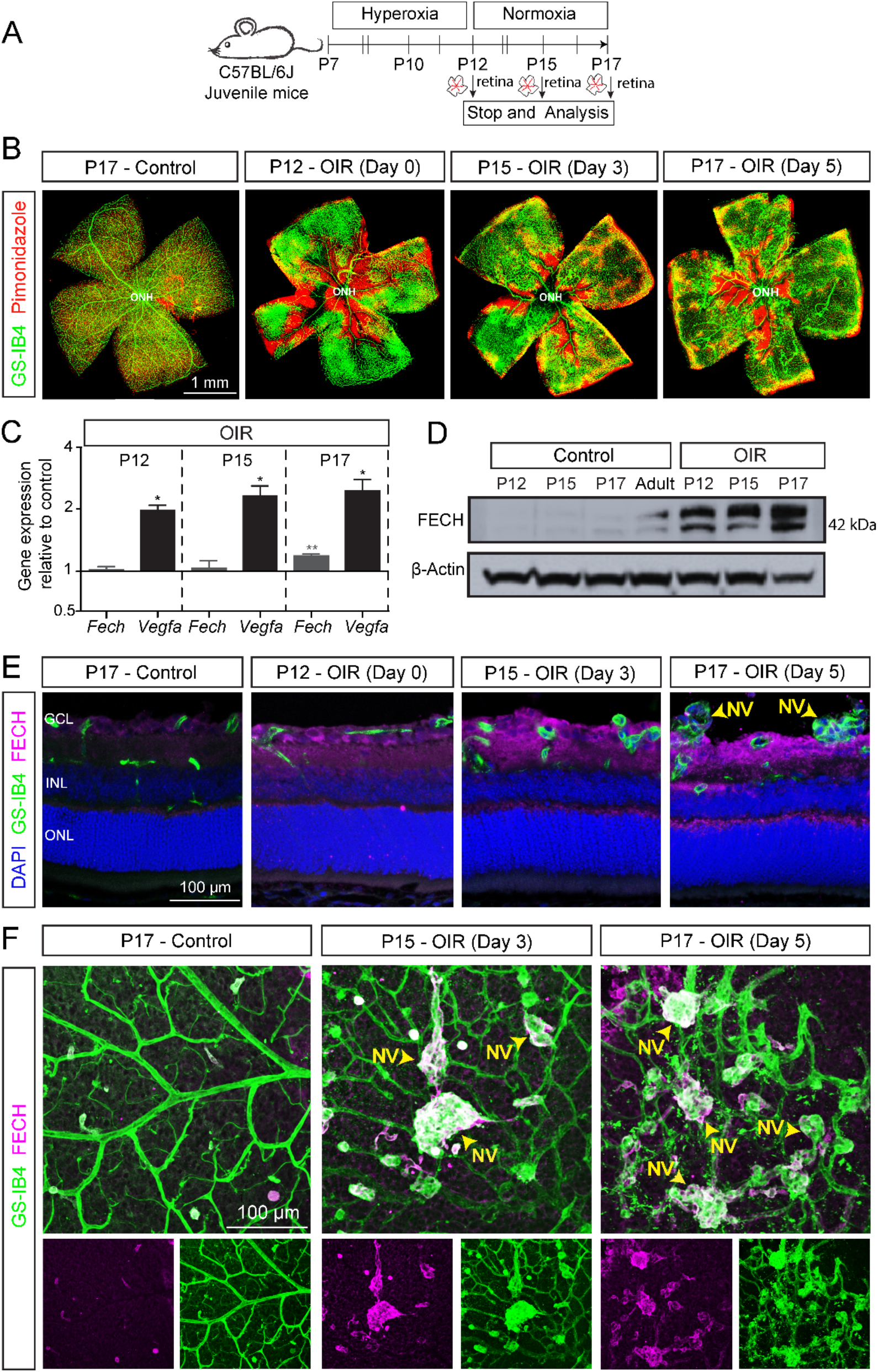
Ferrochelatase (FECH) expression and vascular pathology in the oxygen-induced retinopathy (OIR) mouse model. *A*) Scheme: Juvenile C57BL/6J mice at postnatal day (P) 7 were exposed to hyperoxia chamber for 5 days and returned to room air (normoxia) at P12. Retinal analysis was performed at various time points as indicated. Retinas were isolated for RNA analysis, or fixed, cryosectioned, and immunostained, and retinal sections or whole flatmounts were analyzed. *B*) Retinal flatmount from OIR and control retinas stained with GS-IB4 (green; labels vasculature) and pimonidazole (red; labels hypoxic regions) showing the temporal pattern of hypoxia and formation of pathological neovascular tufts in OIR retina. *C*) Gene expression analysis of *Fech* and *Vegfa* by qPCR of RNA derived from whole retinal samples. Temporal analysis indicated *Fech* and *Vegfa* are upregulated in OIR. C_t_valves from gene expression data were normalized to the housekeeping genes (*Tbp* and *Hprt*) and respective age matched untouched retina control (ΔΔC_t_method). All qPCR reactions were run in technical triplicate, N=3 biological replicates. Data presented as mean ± SEM, *p < 0.05; **p < 0.01 vs. control with Student’s t-test (unpaired, two-tailed). *D*) Immunoblot showing the temporal FECH protein expression in retina from OIR mouse eyes compared to littermate and adult controls; β-actin is a loading control. Pooled retina from three independent animals per condition. *E*) Representative confocal images of retinal sections immunostained for FECH (magenta) and GS-IB4 (green; labels vasculature) show the temporal and spatial pattern of FECH protein expression in the OIR retina. *F*) Representative immunostained enface retinal flatmount images from OIR mice showing upregulation of FECH (magenta) in neovascular tufts stained for GS-IB4 (green; labels vasculature) at P15 and P17 compared to the control. NV: Neovascular tufts; ONH, optic nerve head.

Next, we examined FECH protein expression in the OIR retina by immunoblotting and immunostaining analysis. Temporal analysis showed a significant upregulation of FECH in OIR compared to the age-matched control (Fig. 1*D*). Immunostaining showed FECH expression throughout the retinal layers of the OIR mouse retina compared to age-matched and IgG controls (Fig. 1*E* and Supplemental Fig. S2). Interestingly, enface views of retinal flat mounts showed that FECH is highly colocalized with the neovascular tufts specifically in the superficial vascular plexus at P15 and P17 in OIR compared to control (Fig. 1F and Supplemental Fig. S3). Strikingly, the FECH expression did not overlap with the pimonidazole-adduct hypoxic staining (Supplemental Fig. S4). Thus, our data show that FECH is highly expressed in pathological angiogenic tufts in the murine OIR retina, suggesting FECH is among the mediators of pathological retinal angiogenesis.

### *Fech* ^m1Pas^ mutant abrogates pathological retinal angiogenesis in OIR

To investigate whether FECH directly intervenes in the retinal vascular pathology, we performed OIR in the partial loss-of-function *Fech*^m1Pas^ (ferrochelatase deficiency, mutation 1, Institut Pasteur (36)) murine model (**Fig. 2*A***). Vasculature staining in heterozygous and homozygous *Fech*^m1Pas^ retina from OIR showed substantial reduction in neovascular areas and avascular central regions compared to age-matched wild type control (Fig.2*B*). Retinal neovascularization and vasoobliteration were analyzed quantitatively as described previously (48) as shown in the analysis scheme (Supplemental Fig. S5). Our data revealed a significant reduction of neovascularization (Fig. 2*C*) and vasoobliteration (Fig. 2*D*) in both heterozygous and homozygous *Fech*^m1Pas^ OIR retina compared to wild type control.

**Figure 2.**
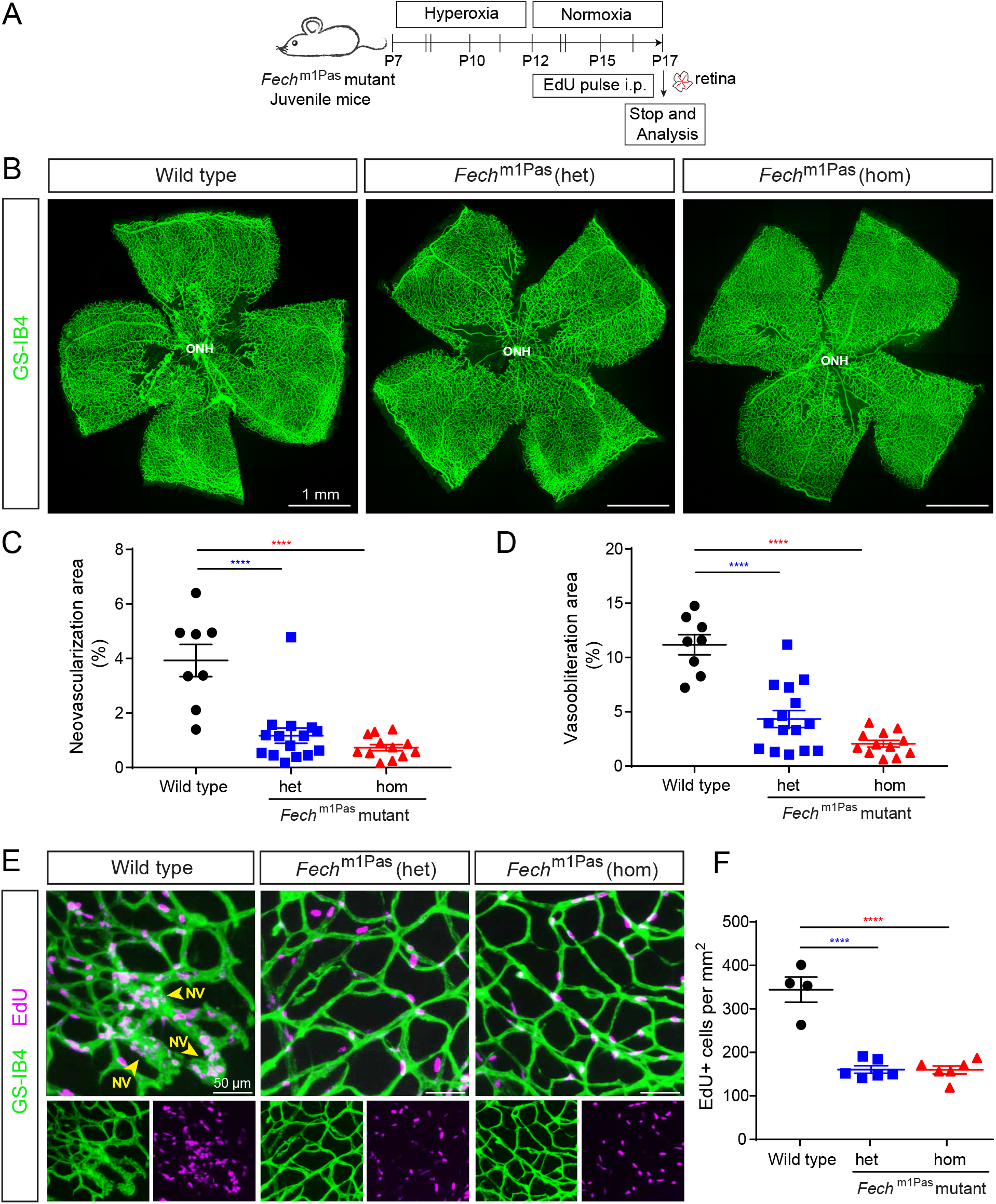
*Fech* mutation abrogates the retinal vascular pathology and proliferation in the OIR mouse model. *A*) Scheme: Partial loss-of-function point mutation *Fech*^m1Pas^ mutant mice at postnatal day (P) 7 were exposed to hyperoxia chamber for 5 days and returned to room air (normoxia) at P12. EdU was administered i.p. at the indicated time points to monitor vascular cell proliferation and retinal analysis was performed at P17 as indicated. *B*) *Fech*^m1Pas^ can ameliorate the two pathological features of the OIR model: neovascularization and vaso-obliteration. P17 OIR retinal flatmount from *Fech*^m1Pas^ mutant stained with GS-IB4 (green; labels vasculature) showing both heterozygous (het; < 50% FECH activity) and homozygous (hom; <10% FECH activity) mutant retinas have reduced retinal neovascularization and vaso-obliteration. *C-D*) Quantification of retinal vaso-obliteration area and retinal neovascularization area in *Fech*^m1Pas^ mutant. Both genotypes have significantly reduced vasoobliteration and neovascularization area compared to wild type controls. Data presented as mean ± SEM, N=8-15, ****p<0.0001 vs. wild type control, one-way ANOVA (Dunnett’s post-hoc test). *E*) Vascular cell proliferation in *Fech*^m1Pas^ mutant retina in OIR mouse model. Retinal flatmount confocal z-stack image stained for GS-IB4 (green, labels vasculature) and EdU (magenta, labels cell proliferation in S-phase) in *Fech*^m1Pas^ mutants suggesting less pathological vascular cell proliferation compared to wild type controls. *F*) Quantifcation of vascular cell proliferation from EdU+ cells per mm^2^ in *Fech*^m1Pas^ mutants and wild type controls. Data presented as mean ± SEM, N=4-6, ****p <0.0001 vs. wild type control, one-way ANOVA (Dunnett’s post-hoc test). NV: Neovascular tufts; ONH, optic nerve head.

Further, we analyzed the vascular cell proliferation in these retinas using EdU staining. *Fech*^m1Pas^ mutant and wild type mice in OIR received EdU (Fig. 2*A*) to identify any cells that entered the cell cycle. Neovascular proliferation was significantly decreased up to 2-fold in both heterozygous and homozygous *Fech*^m1Pas^ mutant compared with wild type control (Fig. 2*E-F* and Supplemental Fig. S6) in OIR. In addition, we tested whether hypoxia is downregulated in *Fech*^m1Pas^ mutants in OIR using pimonidazole-adduct hypoxic staining. Interestingly, we observed a moderate reduction in retinal hypoxia in heterozygous *Fech*^m1Pas^ mutant but a prominent reduction in hypoxia in homozygous *Fech*^m1Pas^ mutant mice compared with wild type mice (Supplemental Fig. S7). Thus, our in vivo data provide evidence that FECH regulates ischemia related pathological retinal angiogenesis.

### Chemical inhibition of FECH blocks retinal angiogenesis and proliferation in OIR

NMPP, a transition-state analogue and selective inhibitor of FECH, is widely used to induce heme deficiency in vitro and ex vivo (53, 54) Our previous studies showed that NMPP inhibited the angiogenic activity in retinal endothelial cells in vitro (23). We hypothesized that inhibition of FECH with NMPP could block the vascular pathology in the OIR retina. We intravitreally injected a single dose of two different concentrations of NMPP (10 and 50 μM, final concentration in the eye) and vehicle at the time of return to normoxia on P12 and quantified the vascular pathology at P17 (**Fig. 3*A***). Both NMPP concentrations significantly reduced the vasoobliteration area and neovascularization area compared to vehicle injected control (Fig. 3*B-D*). Further, we again quantified the vascular cell proliferation using EdU staining. Consistent with our findings with the *Fech* mutant, inhibition of FECH by NMPP treatment drastically diminished the neovascular tuft formation and both concentrations of NMPP significantly reduced vascular cell proliferation (EdU+ cells) up to 2-fold compared to the DMSO vehicle injected control (Fig. 3*E-F* and Supplemental Fig. S8). Thus, our results suggest that chemical inhibition of FECH using NMPP attenuates retinal vascular pathology and improves the revascularization in OIR mice.

**Figure 3.**
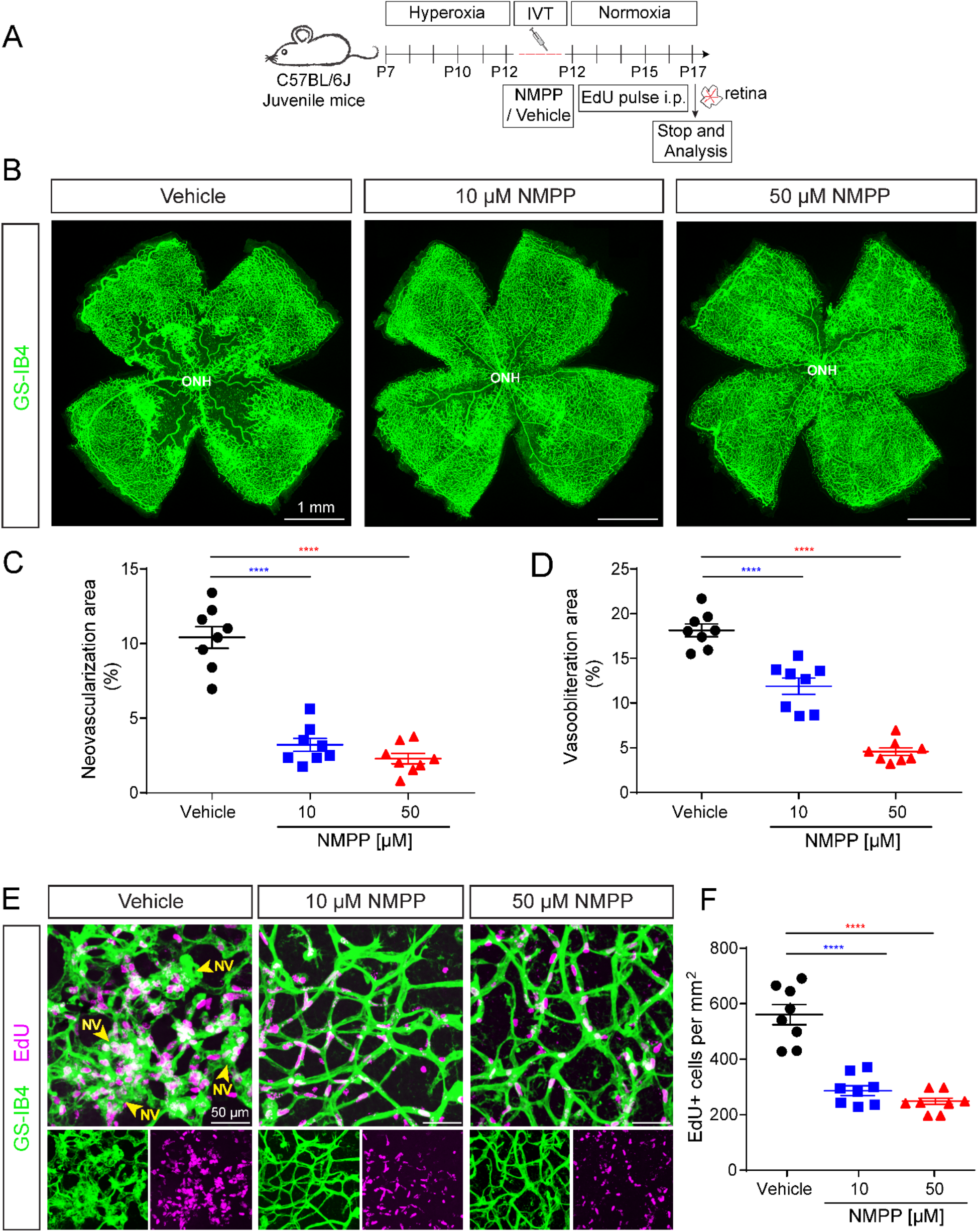
*N*-methylprotoporphyin (NMPP), the canonical FECH small molecule inhibitor, ameliorates retinal angiogensis and cell proliferation in the OIR mouse model. *A*) Scheme: Juvenile C57BL/6J mice at postnatal day (P) 7 were exposed to hyperoxia chamber for 5 days and returned to room air (normoxia) at P12. NMPP was delivered as a single intravitreal injection at P12 in the normoxia phase and EdU was administered i.p. to monitor vascular cell proliferation and retinal analyses were performed at P17 as indicated. *B*) En-face view of P17 OIR retinal flatmouts with GS-IB4 (green; labels vasculature) showing that both NMPP treatments (10 and 50 μM concentration) blocked retinal neovascularization and decreased vaso-obliteration compared to vehicle-injected control. *C-D*) Quantification of retinal neovascularization area and retinal vaso-obliteration area upon vehicle and NMPP treatment. Both NMPP treatments significantly blocked the vasoobliteration and neovascularization area compared to vehicle control. Data presented as mean ± SEM, N=8, ****p<0.0001 vs.vehicle, one-way ANOVA (Dunnett’s post-hoc test). ONH, optic nerve head. *E*) Vascular proliferation in NMPP treated retinas in the OIR mouse model. Retinal flatmount confocal z-stack image stained for GS-IB4 (green, labels vasculature) and EdU (magenta, labels cell proliferation in S-phase) suggesting reduced vascular cell proliferation in NMPP-injected retinas compared to vehicle injected wild-type controls. *F*) Quantifcation of vascular cell proliferation from EdU+ cells per mm^2^ in NMPP (10 and 50 μM) and vehicle controls. Data presented as mean ± SEM, N=8, ****p<0.0001 vs. vehicle, one-way ANOVA (Dunnett’s post-hoc test). NV: Neovascular tufts; ONH, optic nerve head.

### A repurposed anti-fungal drug, griseofulvin, inhibits retinal angiogenesis in OIR

Our previous studies showed that griseofulvin blocks angiogenic activity in retinal endothelial cells in vitro and endothelial sprouting in an ex vivo retinal tissue assay (23). We therefore hypothesized that griseofulvin treatment could diminish neovascularization in the retina in vivo. To this end, we examined the efficacy of griseofulvin in blocking pathological neovascularization in the OIR mouse model. We intravitreally injected two different concentration of griseofulvin (25 and 100 μM, final concentration in the eye) and vehicle at the time of return to normoxia on P12 and quantified the vascular pathology at P17 (**Fig. 4*A***). The higher dose of griseofulvin significantly inhibited pathological retinal neovascularization as compared to vehicle control (Fig. 4*B*). Quantification showed diminished vasoobliteration and neovascularization area with griseofulvin compared to vehicle injected control (Fig. 4*C-D*). Thus, our data demonstrate that griseofulvin not only inhibits pathological neovascularization but also improves retinal revascularization, suggesting a dual effect of vascular repair.

**Figure 4.**
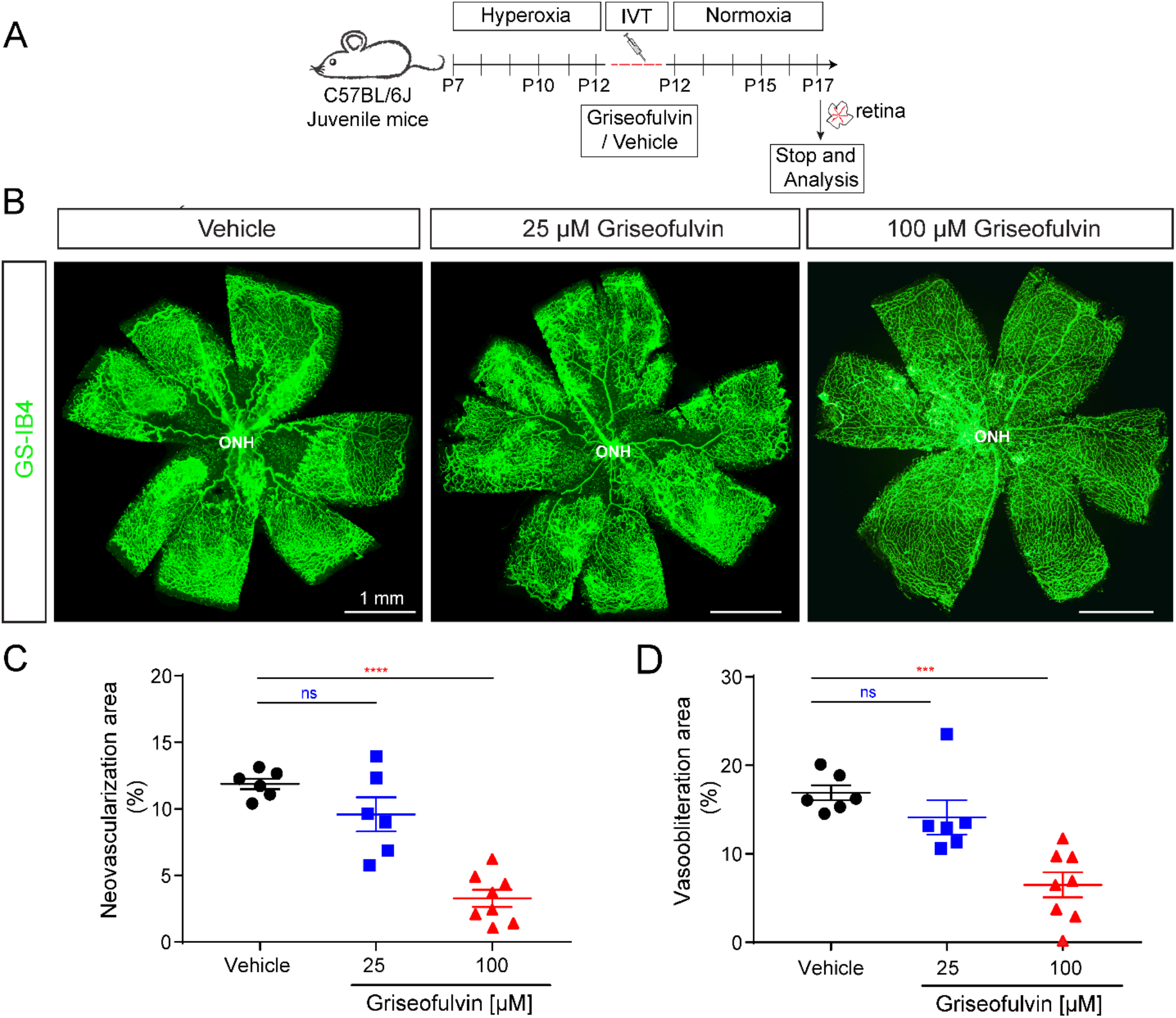
The antifungal drug griseofulvin inhibits retinal angiogenesis and vascular pathology in the OIR mouse model. *A*) Scheme: Juvenile C57BL/6J mice at postnatal day (P) 7 were exposed to hyperoxia chamber for 5 days and returned to room air (normoxia) at P12. Griseofulvin (25 and 100 μM) was delivered as a single intravitreal injection at P12 and retinal analyses were performed at P17 as indicated. *B*) En-face view of P17 OIR retinal flatmouts with GS-IB4 (green; labels vasculature) showing that the higher dose of griseofulvin (100 μM concentration) blocks retinal neovascularization and decreases vaso-obliteration compared to vehicle-injected control. *C-D*) Quantification of retinal neovascularization area and retinal vaso-obliteration area upon griseofulvin and vehicle treatment. 100 μM griseofulvin treatment significantly reduced both neovascularization and vaso-obliteration compared to vehicle injected control. Data presented as mean ± SEM, N=6-8, ns: non significant, ****p<0.0001 vs. DMSO vehicle, one-way ANOVA (Dunnett’s post-hoc test).

### Griseofulvin treatment does not disrupt the retinal structure and retinal functionality in adult mice

Given the potent anti-angiogenic effect of griseofulvin we observed in retinal neovascularization, and the potential for rapid translation of this repurposed drug to human neovascular eye disease therapy, we investigated whether griseofulvin would affect the retinal morphology, vasculature, and retinal function in vivo. We intravitreally injected griseofulvin in normal adult mouse eyes and longitudinally examined the retinal structure using non-invasive OCT, assessed any abnormalities in retinal functionality using ERG, and determined vascular leakage using FA as shown (**Fig. 5*A***). Longitudinal OCT imaging at 5 days after injection (DAI 5) and DAI 14 showed no change in retinal structure and thickness upon griseofulvin treatment compared with vehicle and untouched controls (Fig. 5*B*). Retinal fundus and FA images at DAI 14 showed no aberrant changes in retinal morphology (Fig. 5*C*) and no vascular leakage or disruption of normal vasculature (Fig. 5*D*) upon griseofulvin treatment compared to vehicle injected controls. Scotopic flash ERGs (rod-driven response) were recorded in griseofulvin (100 μM) and vehicle injected animals. Retinal responses in griseofulvin injected animals were comparable to vehicle controls (Fig. 5*E*). Cone-driven photopic ERG showed no differential changes between griseofulvin and vehicle injected animals (Fig. 5*E*). Quantitative analysis confirmed that scotopic a- and b-waves and photopic b-waves were not significantly different between griseofulvin treated animals and vehicle control animals (Fig. 5*F*). These results suggest that griseofulvin treatment does not disrupt the retinal structure and vasculature and does not interfere with retinal function.

**Figure 5.**
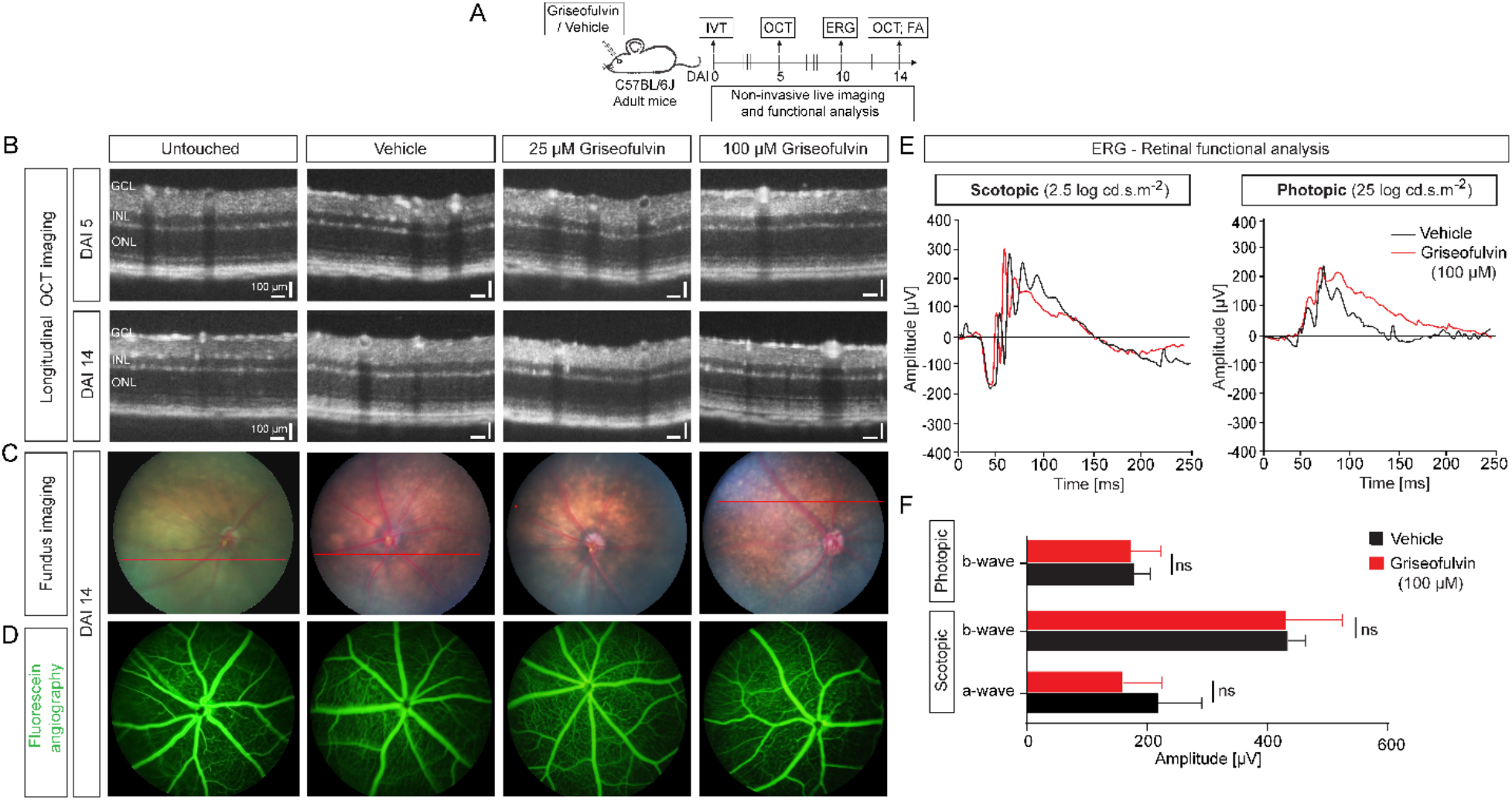
No structural and retinal functional deficits upon griseofulvin treatments in adult mouse eyes. *A*) Scheme: Griseofulvin and vehicle were intravitreally injected in adult C57BL/6J mice. Optical coherence tomography (OCT) was performed at 5 days after injection (DAI 5) and DAI 14 to assess structural changes. Retinal functional analysis was performed using electroretinogram (ERG) at DAI 10. Vascular leakage or changes were assessed by fluorescein angiography (FA) at DAI 14 as indicated. *B*) Longitudinal OCT imaging of vehicle- and griseofulvin (25 and 100 μM)-treated groups at DAI 5 and 14 showed retinal structure is normal. GCL: Ganglion cell layer, INL: Inner nuclear layer, ONL: Outer nuclear layer. *C*) Fundus photographs and *D*) FA images of vehicle and griseofulvin injected groups at DAI 14 showed normal fundus and no vascular leakage. *E*) Retinal functional analysis using ERG. Representative ERG traces of vehicle (DMSO) injected (black) and 100 μM griseofulvin injected (red) eyes, in scotopic (2.5 log cd.s.m^−2^), and photopic (25 log cd.s.m^−2^) conditions at DAI 10, showing no changes under either conditon. *F*) Quantification of the amplitude in both phototopic and scotopic modes showing no significant changes in either a- or b-waves between griseofulvin and vehicle. Data presented as mean ± SD, N=6-10, ns: non-significant, Student’s t-test (unpaired, two-tailed).

### Griseofulvin treatment does not cause retinal cellular toxicity in adult mice

Given that griseofulvin treatment did not change the retinal structure and function, we then explored the morphology and cellular toxicity using immunohistochemistry and histology, respectively, at different time points as shown (**Fig. 6*A***). Hematoxylin and eosin (H & E) stained histological sections of eye samples revealed no evidence of retinal detachment or retinal tears, retinal scarring, or retinal hemorrhage in any of the griseofulvin-injected and control eyes at DAI 7 and 14 (Fig. 6*B*, Supplemental Fig. S9). Quantification of retinal thickness showed the absence of retinal nuclear loss upon griseofulvin treatments compared to vehicle-treated eyes at DAI 7 and 14 (Fig. 6*C*). Further, quantification of the numbers of counted neurons in individual retinal layers showed no significant difference between the griseofulvin-treated eyes and the control groups at DAI 7 and 14 (Fig. 6*D*). These findings suggested that intravitreal injection of griseofulvin did not induce a loss of retinal neurons in adult mice.

**Figure 6.**
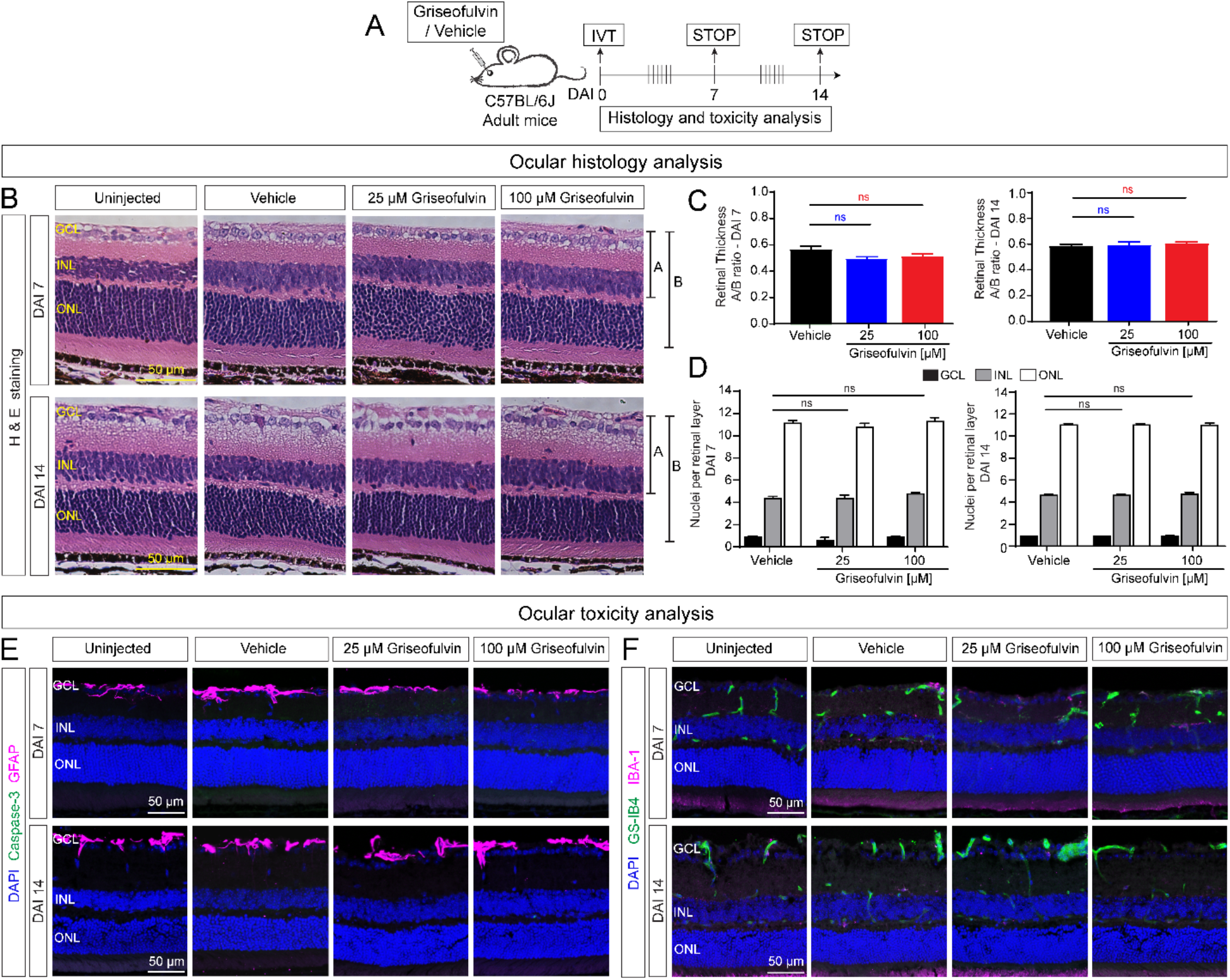
Intravitreal injection of griseofulvin does not cause morphological changes and cellular toxicity in adult mouse eyes. *A*) Scheme: Griseofulvin or vehicle were injected intravitreally in adult C57BL/6J mice. Histology and cytotoxicity analyses were performed at 7 days after injection (DAI 7) and DAI 14 to assess morphological changes. *B*) Representative images of H & E staining from uninjected control, vehicle, and griseofulvin treatments. Vehicle- and griseofulvin (25 and 100 μM)-treated groups at DAI 7 and 14 showed normal retinal morphology and retinal thickness in all the layers. *C*) Quantifiation of the retinal thickness of the indicated A/B ratio showed no significant changes in retinal layers at DAI 7 and DAI 14 upon griseofulvin treatments (25 and 100 μM concentration) compared to vehicle-injected control. *D*) Quantification of the retinal layer thickness in terms of cell nuclei per nuclear layer showed no significant change in any retinal nuclear layer at DAI 7 and DAI 14 upon griseofulvin treatments (25 and 100 μM concentration) compared to vehicle-injected control. Data presented as mean ± SEM, N=6-10, ns: non-significant, griseofulvin treatment vs DMSO vehicle, one-way ANOVA (Dunnett’s post-hoc test). *E*) Representative confocal images of retinal sections from vehicle and griseofulvin treated groups, immunostained for caspase 3 (green) and GFAP (magenta) at DAI 7 and 14 revealed no differences between griseofulvin treatment and vehicle control in GFAP and cleaved caspase 3 as markers of retinal gliosis and apoptosis respectively. *F*) Immunofluorescence images show GS-IB4 (green; labels vasculature) and IBA-1 (magenta; labels microglia) staining in vehicle and griseofulvin treated retinal sections at DAI 7 and 14. Griseofulvin treatment did not disrupt the normal vasculature and did not induce inflammation via microglia activation. GCL: Ganglion cell layer, INL: Inner nuclear layer, ONL: Outer nuclear layer.

To further test the cellular toxicity of griseofulvin besides retinal morphology, retinal sections were examined for cell death, stress and inflammation markers. Retinal sections were stained for the apoptotic cell death (cleaved active caspase 3) and the reactive gliosis or stress marker, glial fibrillary acidic protein (GFAP) and in griseofulvin- and vehicle-treated eyes. Neither at DAI 7 nor DAI 14 was there an increase in GFAP or caspase 3 expression in griseofulvin-treated eyes compared to vehicle-treated eyes (Fig. 6*E*, Supplemental Fig. S10). Finally, retinal sections were stained for microglia (IBA-1, marker for inflammation) and GS-IB4 (marker for vasculature). DAI 7 and 14 immunostaining revealed that there was no activation of microglia due to injury or inflammation and the retinal vascular plexus appeared quite normal between the griseofulvin-treated eyes and vehicle-treated eyes (Fig. 6*F*, Supplemental Fig. S11). Thus, our data suggest the absence of ocular toxicity of griseofulvin on adult retina.

## DISCUSSION

The association of neovascularization with the most common causes of blindness has encouraged the search for a better understanding of how angiogenesis is regulated in health and disease (10, 29, 55). Targeting VEGF and other proangiogenic factors has been successful in blocking pathological neovascularization but challenges persist to devise approaches to revascularize the ischemic retina under pathological conditions, and avoid the risks of anti-VEGF therapies, especially in neonates. In this study, we provide evidence that FECH upregulation in vasculature contributes to pathological retinal neovascularization; inhibiting FECH either by genetic or chemical approaches offers potential as a novel treatment for ischemic proliferative vascular retinopathies (Supplemental Fig. S12).

Previously, we have shown that FECH is upregulated in human wet AMD patient and murine laser-induced choroidal neovascularization eyes (23). In OIR, we observed FECH overexpression in retina, notably at the protein level but also more modestly at the mRNA level. We thought this upregulation was likely due to hypoxia, since hypoxia upregulates the key enzymes responsible for heme biosynthesis (56) and notably, FECH expression was upregulated in Hep-G2, K562, and HEL cells during hypoxia via induction of hypoxia inducible factor-1 (HIF-1) (57). Intriguingly, FECH expression was co-localized with epiretinal neovascular tufts and widely distributed across the inner nuclear layer and plexiform layers in the mouse OIR retina. However, FECH rarely co-localized with the pimonidazole-adduct hypoxia staining, suggesting that other factors drive FECH expression.

The *Fech*^m1Pas^ mouse is a widely used in vivo model to study erythropoietic protoporphyria (EPP), a human disease characterized by skin photosensitivity and partial FECH deficiency that results in excessive accumulation of photoreactive PPIX in blood cells and other organs (58). The *Fech*^m1Pas^ allele encodes an M98K missense mutation that reduces *Fech* heme synthesis activity by ~50% in heterozygotes and ~95% in homozygotes (36, 59). In our genetic approach, mice with mutated *Fech* exhibited reduced pathological retinal neovascularization. In fact, *Fech*^m1Pas^ mutants had improved vascularization associated with ischemia and intriguingly, in homozygous *Fech*^m1Pas^ mutants, OIR-driven hypoxia was reduced. In our chemical approach, we saw that in the eye, NMPP treatment blocked pathological retinal neovascularization, consistent with our findings in the *Fech* mutant. In agreement, NMPP treatments blocked pathological epiretinal cell proliferation (EdU+ cells). A decrease in vasoobliteration area (capillary drop out) during hypoxia is a key characteristic feature of physiological retinal revascularization. Interestingly, NMPP treatment reduced vasoobliteration area, suggesting that FECH inhibition improves the physiological revascularization of the central retina in the ischemia driven OIR model. We believe this is the first report to demonstrate that inhibiting FECH not only reduces neovascularization but promotes vascular repair and protection in ischemic retinopathy. We speculate that loss of FECH might enable appropriate angiogenic signals for revascularization, by reduction in inflammation (60) and enhanced astrocyte-endothelial interactions for physiological vascular repair (61). FECH is thus an important regulator for retinal angiogenesis in pathological conditions associated with injury or stress.

Our prior studies in choroidal neovascularization showed that griseofulvin blocked neovascularization in this vascular bed, comparable with anti-VEGF treatment in adult mice (23). In OIR as studied here in juvenile mice, intravitreal griseofulvin treatment blocked retinal neovascularization and, strikingly, stimulated the revascularization of the central part of the retina and improved physiological angiogenesis, comparable to NMPP treatment. Intravitreal injections of griseofulvin did not change retinal structure and morphology, with no vascular abnormalities. These findings concur with the phenotype of patients with EPP, who despite having constitutively reduced FECH, primarily show skin photosensitivity and (rarely) liver damage (58), with only one documented case of a (possibly incidental) ocular phenotype (62). In griseofulvin-treated eyes, retinal function was normal with no aberrant cell death, microglia activation or gliosis, suggesting no cellular toxicity or stress. Hence, griseofulvin could be a potential candidate for treatment of retinal neovascular disease, perhaps as intravitreal injections or topical formulations, likely with minimal systemic side effects. An effective concentration of griseofulvin in the neonatal human eye should be achievable using a standard 25 μL injection. However, further long-term ocular toxicity studies are indicated.

The exact mechanisms of how FECH loss modulates angiogenesis remain to be determined, but there is growing evidence that FECH, via heme, acts as a master regulator of several angiogenic pathways. Heme, as an iron containing porphyrin cofactor, is crucial for several biological processes such as erythroid differentiation, oxygen transport, cellular signaling, and drug metabolism (63, 64). Blocking vascular heme synthesis by inhibition of FECH using NMPP downregulates endothelial nitric oxide synthase (eNOS) and soluble guanylyl cyclase mediated vascular responses in vitro (23, 54). Heme oxygenase (HO-1) is an essential enzyme responsible for degrading heme into ferrous iron, biliverdin and carbon monoxide, and is involved in formation of blood vessels (65) and preventing vascular inflammation (66). Heme metabolites contribute to angiogenesis after chronic injury and induction of HO-1 with hemin in endothelial cells resulted in enhanced carbon monoxide production and elevated VEGF synthesis (67). VEGF regulates HO-1 expression in endothelial cells and inhibition of HO-1 reduced VEGF-induced angiogenesis in endothelial cells (65, 68). Previously, we have shown that FECH depletion decreases eNOS, HIF-1 and VEGF receptor 2 levels in retinal endothelial cells (23). Further, supplementation with hemin restores these, suggesting the essential interaction of heme and other proangiogenic regulators.

Intriguingly, recent work revealed that heme production was reduced in endothelial cells upon deletion of phosphoglycerate dehydrogenase (PHGDH), a key enzyme of the serine synthesis pathway and addition of hemin under PHGDH knockdown conditions rescued the angiogenic defects. In addition, endothelial specific PHGDH deletion led to vascular defects with decreased glutathione and NADPH synthesis in vivo (69), suggesting serine synthesis is essential for heme production in endothelial cells. Further, heme depletion selectively reduced the protein expression and activity of components of the mitochondrial electron transport chain, which could lead to enhanced mitochondrial dysfunction in endothelial cells (69–71). It will be interesting to ascertain the downstream interactions of hemoproteins upon FECH depletion in retinal and choroidal angiogenesis mouse models.

In conclusion, mutation of *Fech* and chemical inhibition reduces pathological retinal neovascularization and promotes physiological angiogenesis, suggesting a dual effect on vascular repair upon FECH inhibition. The anti-fungal drug griseofulvin, metabolized into a FECH inhibitor, is already FDA approved as a systemic agent and has promising safety data in the eye. Hence, there is strong potential to repurpose griseofulvin for the treatment of ischemia-related retinopathies and other neovascular ocular diseases.

## Abbreviations

BSA: bovine serum albumin
DAI: days after injection
DMSO: dimethyl sulfoxide
EdU: 5-ethynyl-2′-deoxyuridine (EdU)
ERG: electroretinogram
eNOS: endothelial nitric oxide synthase
EPP: erythropoietic protoporphyria
ETC: electron transport chain
FA: fluorescein angiography
FECH: ferrochelatase
GCL: ganglion cell layer
GFAP: glial fibrillary acidic protein
GS-IB4: *Griffonia simplicifolia* isolectin B4
H&E: hematoxylin and eosin
HIF-1: hypoxia inducible factor 1
HO-1: heme oxygenase 1
INL: inner nuclear layer
NMPP: *N*-methyl protoporphyrin
OCT: optical coherence tomography
OIR: oxygen induced retinopathy
ONL: outer nuclear layer
PBS: phosphate buffered saline
PDR: proliferative diabetic retinopathy
PFA: paraformaldehyde
PPIX: protoporphyrin IX
ROP: retinopathy of prematurity
VEGF: vascular endothelial growth factor

## ACKNOWLEDGMENTS

We thank members of the Corson Laboratory for comments on the manuscript; Ms. Kamakshi Sishtla for her valuable technical help in mouse genotyping; Dr. Ashay Bhatwadekar, IUSM for helpful comments on the manuscript; Dr. Keith W. Condon, IUSM Histology core for retinal histology; and members of Dr. Harikrishna Nakshatri’s Laboratory for technical assistance in preparing retinal cryosections. This study was supported by NIH/NEI grant R01EY025641 and BrightFocus Foundation Macular Degeneration Research grant M2019069. Conflict of Interest Statement: S. Pran Babu and T. W. Corson are named inventors on patent applications related to this work.

## AUTHOR CONTRIBUTIONS

S. Pran Babu and T. W. Corson designed research; S. Pran Babu and D. White performed research; S. Pran Babu, D. White, and T. W. Corson analyzed data; and S. Pran Babu and T. W. Corson wrote the paper.

