## Supplemental Figures for "Ferrochelatase regulates retinal neovascularization"

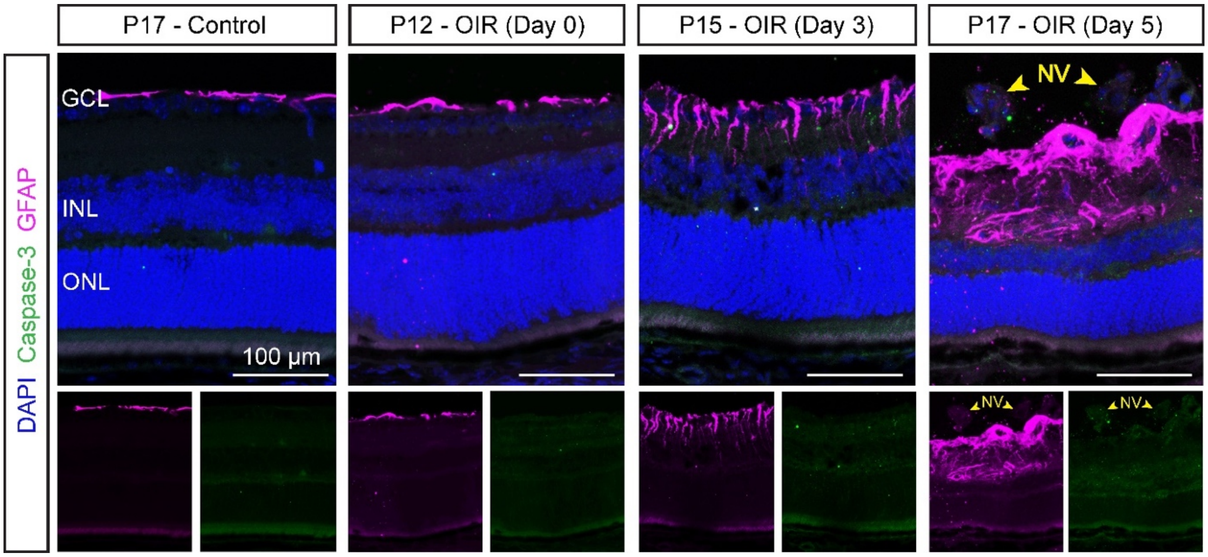


**Supplemental Figure S1.** Temporal pattern of Caspase-3 and GFAP expression in the OIR mouse model. Immunostained retinal sections showing the upregulation of gliosis marker, GFAP (magenta) and cell death marker, caspase 3 (green) at P15 and P17 in the GCL and INL of OIR eyes. GCL: Ganglion cell layer, INL: Inner nuclear layer, ONL: Outer nuclear layer, NV: Neovascular tufts.

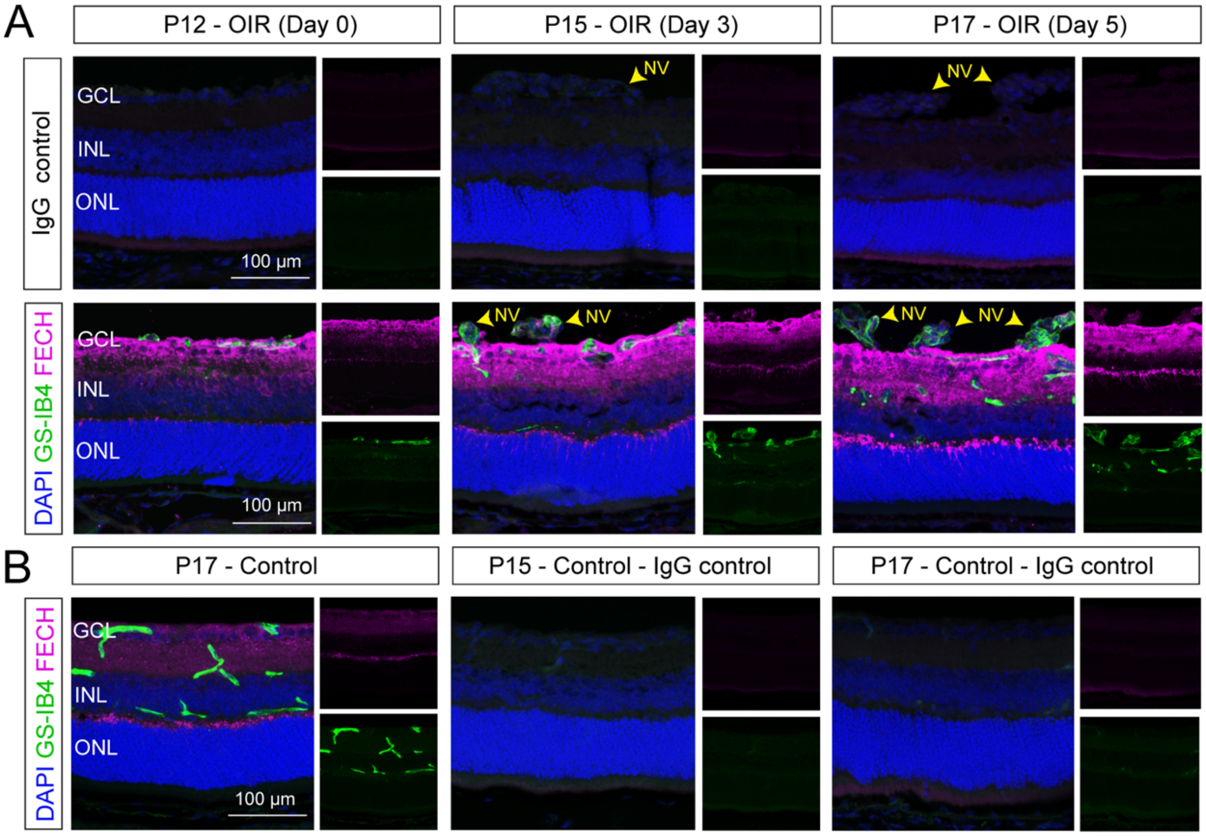


**Supplemental Figure S2.** Temporal analysis of vasculature and FECH expression *A*) in the OIR mouse model and *B*) in control (normoxic) animals. Immunostained retinal sections showing the upregulation of FECH (magenta) and vascular pathology defects as GS-IB4 (green; labels vasculature) at P15 and P17 in the GCL and INL of OIR eyes. GCL: Ganglion cell layer, INL: Inner nuclear layer, ONL: Outer nuclear layer, NV: Neovascular tufts.

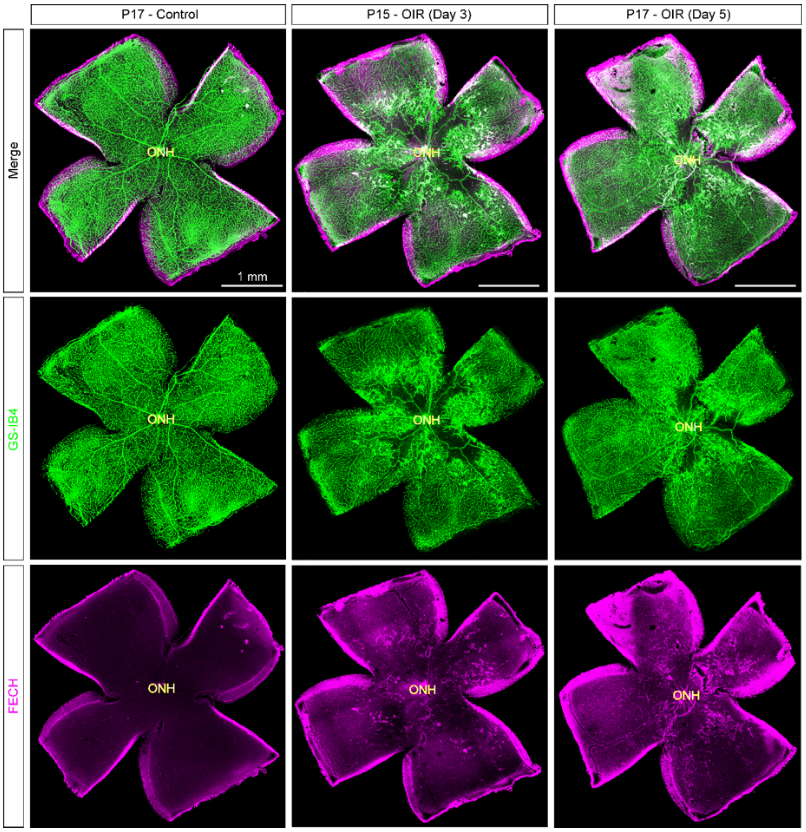


**Supplemental Figure S3.** Temporal expression of FECH and vascular pathology in the OIR mouse model. Enface overview of retinal flatmounts from OIR retina immunostained with GS-IB4 (green; labels vasculature) and FECH (magenta) in P17 control, P15, and P17 OIR retinas. FECH upregulation is seen over the OIR retinas and specifically in the central area of the retinas where the neovascularization occurs, compared to the control (FECH signal along cut edges is artifactual). ONH, optic nerve head.

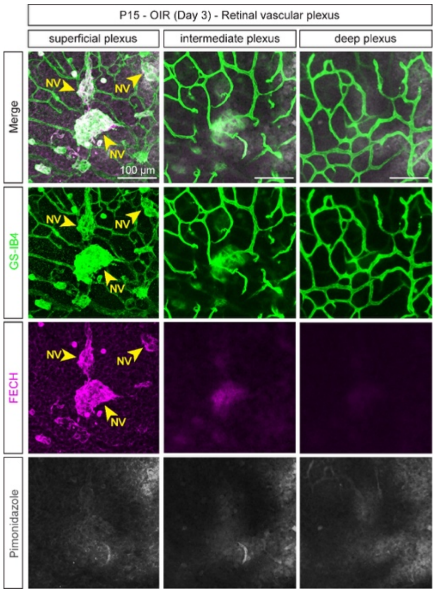


**Supplemental Figure S4.** Assessment of FECH and hypoxia co-localization in the retinal vascular plexus in the OIR model. Confocal z-stack images of retinal flatmounts stained for GS-IB4 (green, labels vasculature), pimonidazole adducts (gray; labels hypoxic regions) and FECH (magenta) from P15 OIR mouse retina. Depth projections show that FECH is colocalized with the pathological vascular tufts in the superficial vascular plexus but not in the intermediate and deep vascular plexus, and FECH is not colocalized with pimonidazole adducts in any layer of the retinal vasculature. NV: Neovascular tufts.

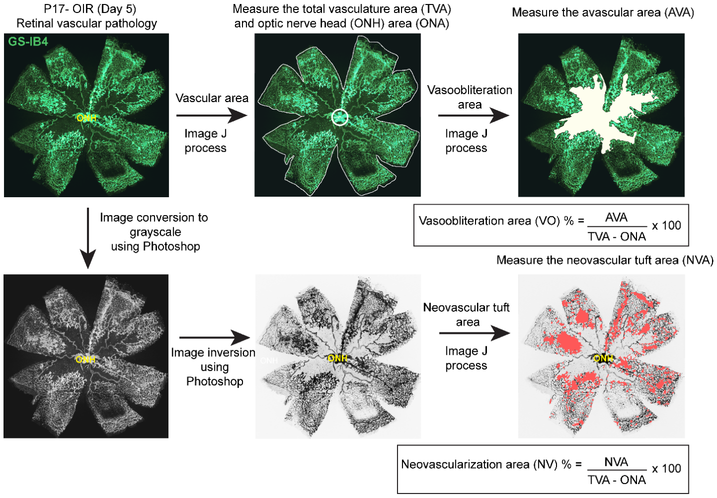


**Supplemental Figure S5.** Schematic workflow illustration of quantification of vasoobliteration (VO) and neovascularization (NV) area in flatmounts of retinas with OIR. Representative image of retinal vasculature from wild type animal with induced OIR at P17, stained with GS-IB4 (green, labels vasculature). Areas of vasoobliteration and pathological neovascularization were highlighted in white and coral respectively. Quantification of retinal vasoobliteration area (VO) and neovascularization area (NV) in OIR is represented as percent of total retinal vasculature area (TVA) excluding optic nerve head area (ONA). ONH, optic nerve head.

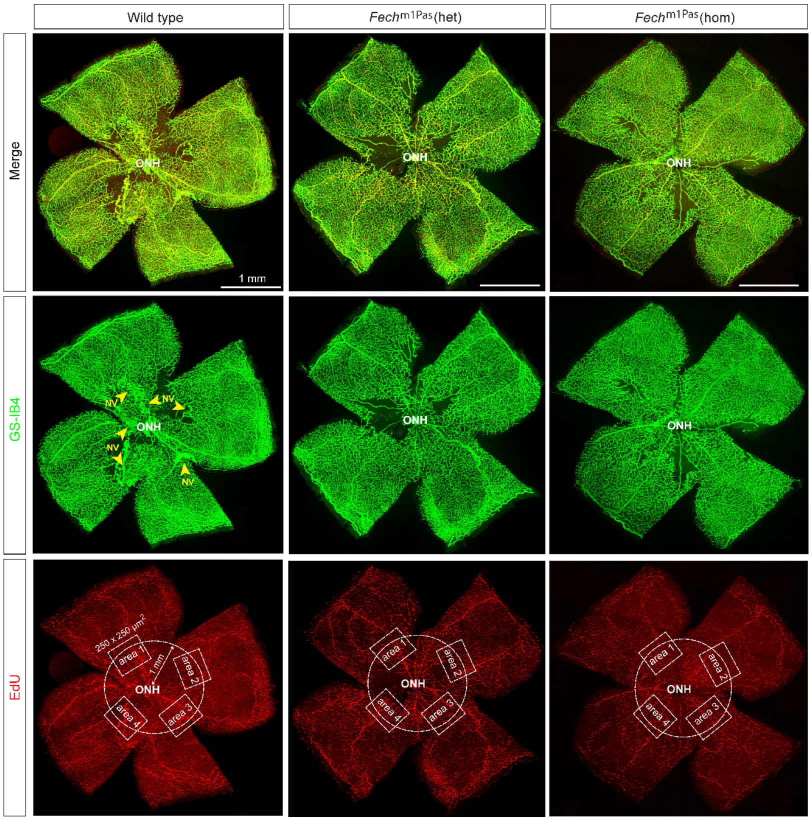


**Supplemental Figure S6.** Vascular cell proliferation and its analysis in *Fech*$^{m1Pas}$ mutant mice OIR retinal flatmounts. Enface retinal flatmount overview images from P17 *Fech*$^{m1Pas}$ mutants with OIR immunostained with GS-IB4 (green, labels vasculature) and EdU (red; labels S-phase in cell cycle). Confocal z-stack imaging was performed over an area of 250 μm$^2$ (inset boxes, area) at 1 μm intervals on retina flatmounts with EdU across the entire sample depth, and z-stack slice images were flattened for analysis. For EdU cell proliferation counts, image areas were selected in the middle of each retinal "petal", along a circle of 1 mm radius from the optic nerve head region as shown by dashed circles. The optic nerve head region as well as the peripheral edges were excluded from the EdU analysis. NV, neovascular tufts; ONH, optic nerve head.

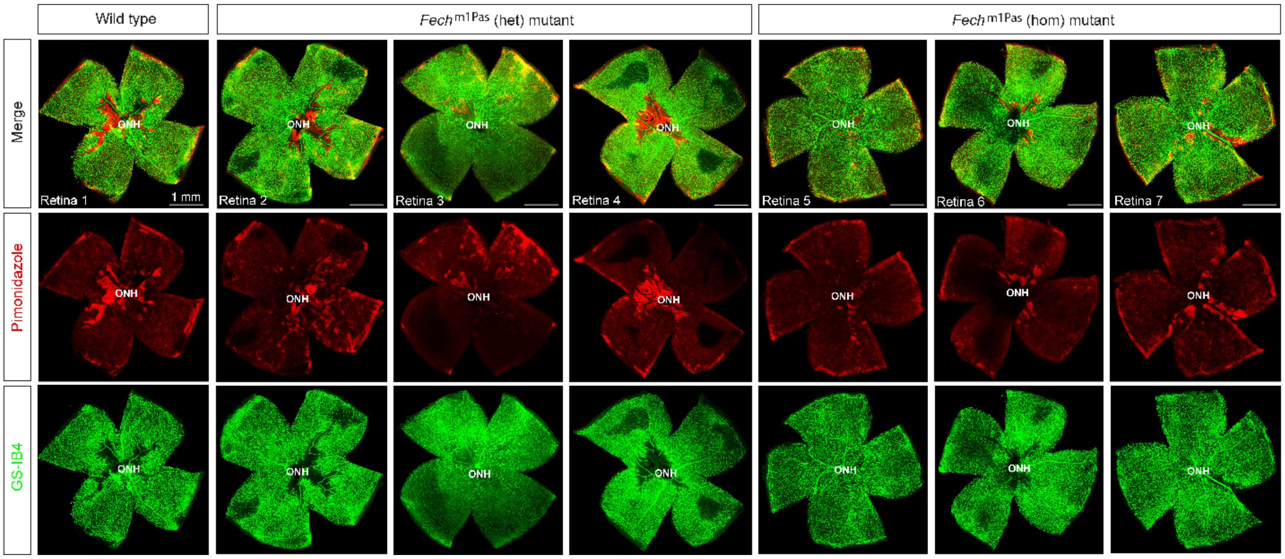


**Supplemental Figure S7.** Differential hypoxia regulation in *Fech*$^{\mathrm{m1Pas}}$ mutants. Enface overview of P17 OIR retinal flatmounts from wild type, heterozygous (het) and homozygous (hom) *Fech*$^{\mathrm{m1Pas}}$ mutants stained for GS-IB4 (green; labels vasculature) and pimonidazole adducts (red; labels hypoxic regions). The hypoxic regions were reduced in *Fech*$^{\mathrm{m1Pas}}$ (hom) mutants compared to wild type and *Fech*$^{\mathrm{m1Pas}}$ (het) mutant retinas. ONH; optic nerve head.

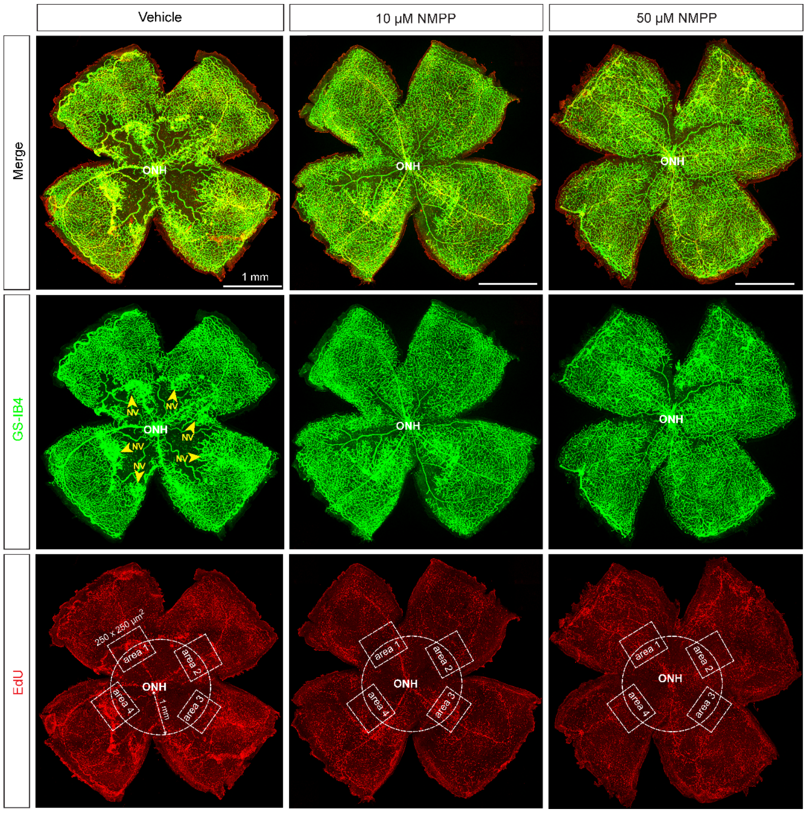


**Supplemental Figure S8.** Vascular cell proliferation and its analysis in NMPP treated mice OIR retinal flatmounts. Enface retinal flatmount overview images from P17 vehicle and NMPP-treated groups with OIR immunostained with GS-IB4 (green, labels vasculature) and EdU (red; labels S-phase in cell cycle). Confocal z-stack imaging was performed over an area of 250 μm$^2$ (inset boxes, area) at 1 μm intervals on retina flatmounts with EdU across the entire sample depth, and z-stack slice images were flattened for analysis. For EdU cell proliferation counts, image areas were selected in the middle of each retinal "petal", along a circle of 1 mm radius from the optic nerve head region as shown by dashed circles. The optic nerve head region as well as the peripheral edges were excluded from the EdU analysis. NV, neovascular tufts; ONH, Optic nerve head.

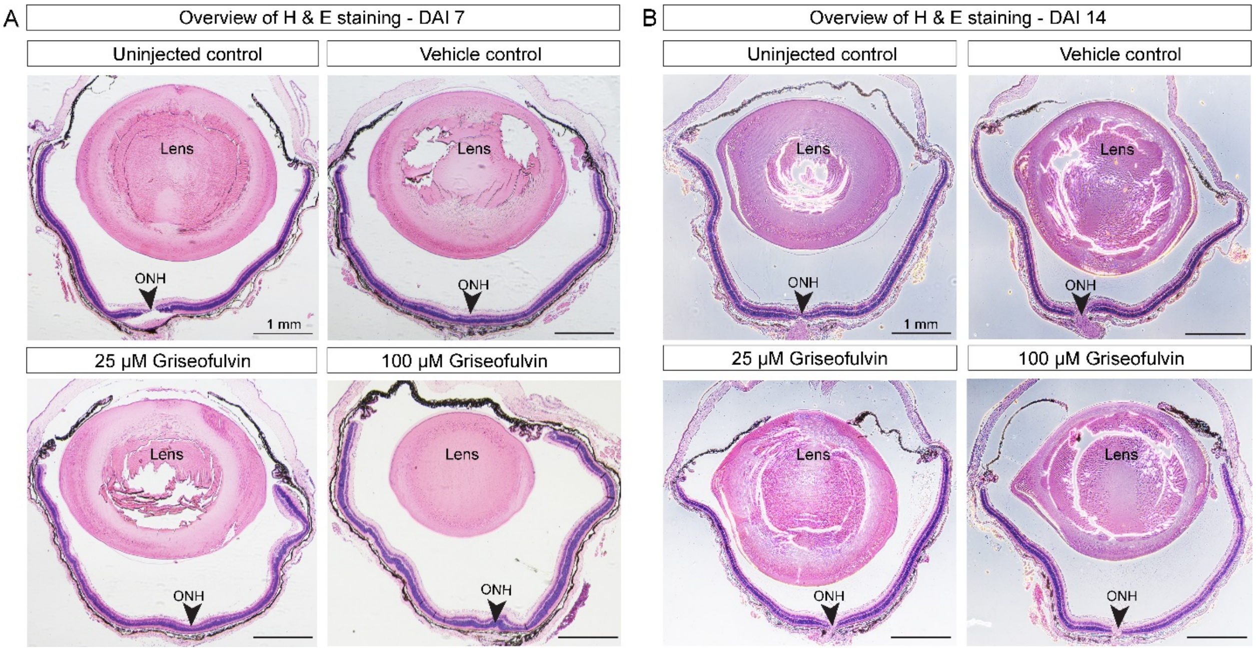


**Supplemental Figure S9.** Histopathology of representative whole mouse eye sections close to the optic nerve-pupillary axis from griseofulvin and vehicle injected controls. Overview of the H & E histological sections at *A*) 7 days after injection (DAI 7) and *B*) DAI 14, reveals no substantial changes in retinal morphology in griseofulvin- and vehicle-treated groups compared to uninjected control. ONH, Optic nerve head.

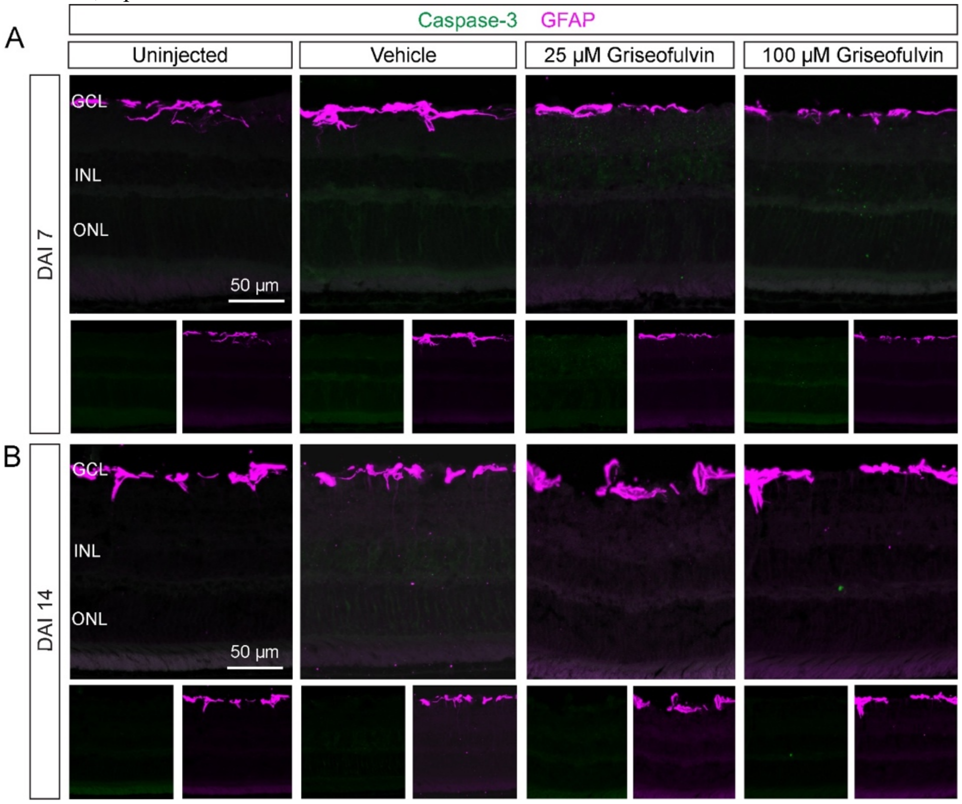


**Supplemental Figure S10.** Caspase-3 and GFAP expression in vehicle- and griseofulvin-treated retinal sections from adult mouse eyes. Individual confocal fluorescence images show apoptosis (Caspase-3; green) and gliosis (GFAP; magenta) signals in vehicle- and griseofulvin- treated retinal sections in adult mouse at *A*) 7 days after injection (DAI 7) and *B*) DAI 14. No aberrant retinal cell death nor reactive gliosis were seen upon griseofulvin and vehicle treatment compared to uninjected control. GCL: Ganglion cell layer, INL: Inner nuclear layer, ONL: Outer nuclear layer.

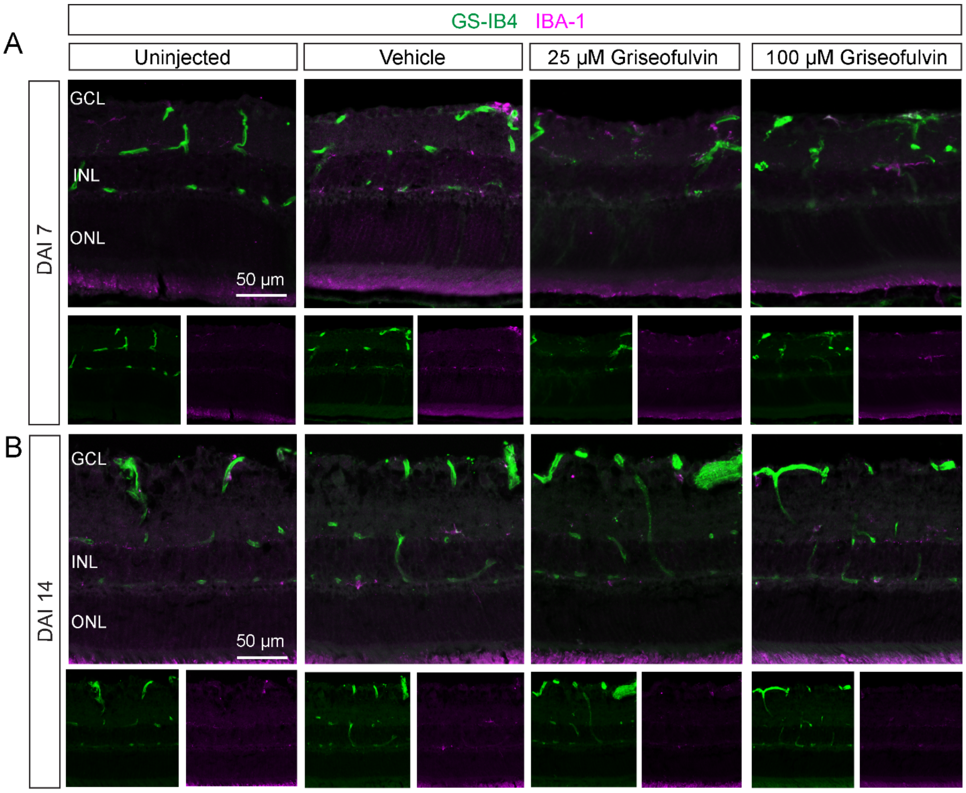


**Supplemental Figure S11.** Microglia activation and vasculature in griseofulvin-treated retinas from adult mouse eyes. Individual confocal fluorescence images show vasculature (GS-IB4; green) and microglia (IBA-1; magenta) in vehicle- and griseofulvin-treated retinal sections in adult mouse at *A*) 7 days after injection (DAI 7) and *B*) DAI 14. No vascular changes nor microglia activation were seen upon griseofulvin and vehicle treatment compared to uninjected control. GCL: Ganglion cell layer, INL: Inner nuclear layer, ONL: Outer nuclear layer.

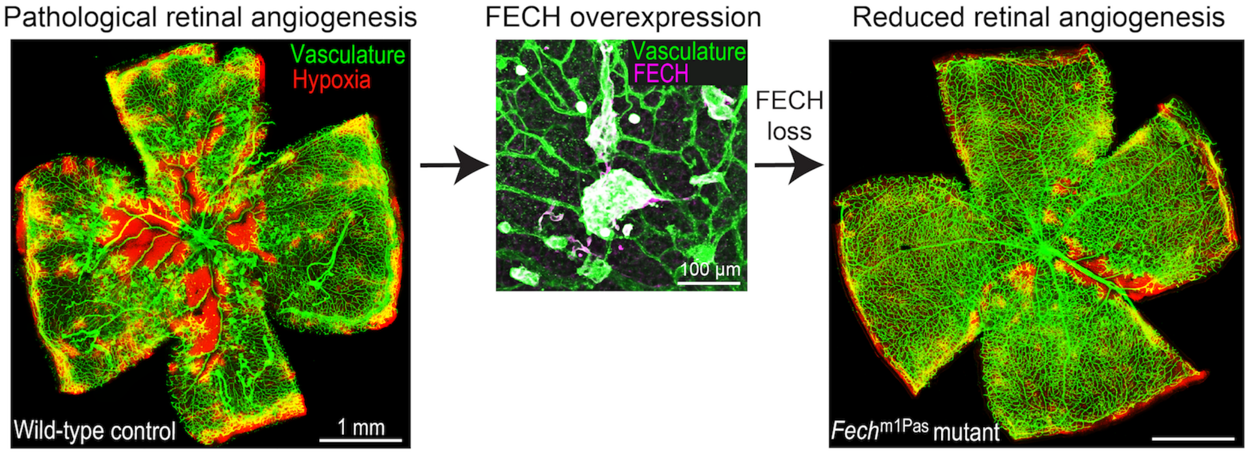


**Supplemental Figure S12.** Summary of findings. In OIR, FECH is overexpressed; loss of FECH by chemical or genetic methods reduced retinal angiogenesis and hypoxic areas.
